# Computational Modeling of the Human Compound Action Potential

**DOI:** 10.1101/2022.08.26.505458

**Authors:** Yousef Alamri, Skyler G. Jennings

## Abstract

The auditory nerve (AN) compound action potential (CAP) is an important tool for assessing auditory disorders and monitoring the health of the auditory periphery during surgical procedures. The CAP has been mathematically conceptualized as the convolution of a unit response (UR) waveform with the firing rate of a population of AN fibers. Here, an approach for predicting experimentally-recorded CAPs in humans is proposed, which involves the use of human-based computational models to simulate AN activity. CAPs elicited by clicks, chirps, and amplitude-modulated carriers were simulated and compared with empirically recorded CAPs from human subjects. In addition, narrowband CAPs derived from noise-masked clicks and tone bursts were simulated. Many morphological, temporal, and spectral aspects of human CAPs were captured by the simulations for all stimuli tested. These findings support the use of model simulations of the human CAP to refine existing human-based models of the auditory periphery, aid in the design and analysis of auditory experiments, and predict the effects of hearing loss, synaptopathy, and other auditory disorders on the human CAP.

## I. INTRODUCTION

Electrocochleography (ECochG) is a technique for recording cochlear and auditory nerve (AN) potentials in response to sound stimulation (Eggermont, 2017). These potentials may be recorded in humans by placing a transtympanic electrode on the cochlear promontory (e.g., Portmann and Aran, 1971), or by placing an extratympanic or tympanic electrode in the ear canal or on the tympanic membrane (TM), respectively (e.g., Pappas *et al.*, 2000; Simpson *et al.*, 2020). In clinical settings, ECochG may serve as a tool for intraoperative monitoring of cochlear insult during cochlear implant insertion (e.g., Campbell *et al.*, 2015), and for the assessment and diagnosis of auditory disorders such as Ménière’s disease (e.g., Ruth and Lambert, 1989) and auditory neuropathy (e.g., Santarelli and Arslan, 2002). A typical ECochG recording includes three overlapping potentials – the cochlear microphonic (CM), summating potential (SP), and compound action potential (CAP). The CM is an alternating current (AC) potential dominated by the electrical activity of the outer hair cells (OHCs) (Dallos, 1972), whereas the SP is a direct current (DC) potential dominated by the electrical activity of OHCs and inner hair cells (IHCs) (Durrant *et al.*, 1998), and may include a neural component (Pappa *et al.*, 2019). The CAP, which is primarily characterized by two brief negative deflections (N_1_ and N_2_), represents the synchronous firing of AN fibers (ANFs) and is typically elicited in response to the onset of a stimulus (Derbyshire and Davis, 1935).

Goldstein and Kiang (1958) proposed a model that captures the relationship between the activity of a single-unit within a population of ANFs and the CAP recorded from an electrode placed on or near the round window. If each neural unit generates a unit response (UR) waveform upon the discharge of an action potential, the CAP measured from the round window can be expressed as

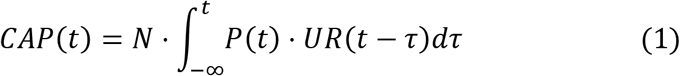

where *N* is the number of ANFs in the population, *P*(*t*) is the sound-evoked probability density function of ANF discharges, and *UR*(*t*) is the unit response waveform resulting from the firing of an action potential. Several simplifying assumptions were made by Goldstein and Kiang (1958) with regard to this convolution-based model. First, each action potential generated by an ANF was assumed to result in the same UR waveform, regardless of the fiber’s characteristic frequency (CF) or spontaneous rate (SR) of discharge. Second, each UR contributed equally to the CAP with no differential weight based on the CF or SR. Finally, Goldstein and Kiang (1958) made the assumption that *P*(*t*) was the same for peripheral and cortical generators.

Several studies have estimated the UR from data measured from laboratory animals and humans using a variety of techniques. For example, acoustical masking of the CAP has been used to estimate the UR in guinea pigs (Teas *et al.*, 1962). Similarly, spike-triggered averaging has been employed to estimate the UR in cats (Kiang *et al.*, 1976) and guinea pigs (Prijs, 1986). Elberling (1976) empirically estimated the human UR associated with an electrode placed on the TM using a masking technique to limit ANF activity to the base of the cochlea. Specifically, the average of CAPs recorded from the 8-20 kHz cochlear region of several human subjects served as an estimate of the UR after correcting for the expected traveling wave delay. The rationale for limiting the recording to these unmasked basal ANFs was founded on the assumption that these fibers fire synchronously and their discharge pattern [P(t)] resembles an impulse and thereby producing a CAP that is largely determined by the shape of the UR.

Several approaches have been employed to estimate the probability density function of ANF discharges, *P*(*t*). For instance, de Boer (1975) used a cochlear model to estimate *P*(*t*) as the output of a probabilistic pulse generator driven by an input signal designed to simulate the cochlear response to a tone burst (TB) stimulus for the auditory system of a cat. Another approach, which is adopted for this study, relied on modeling *P*(*t*) from the post-stimulus time histogram (PSTH) (Kiang *et al.*, 1976; Versnel *et al.*, 1992). For example, Chertoff (2004) modeled *P*(*t*) as the envelope of TB-evoked PSTHs measured from guinea pigs using parameterized gamma functions.

Developments in computational models of the auditory system have enabled the simulation and interpretation of perceptual (e.g., Jennings *et al.*, 2011; Jorgensen and Dau, 2011) and evoked-potential (e.g., Dau, 2003; Verhulst *et al.*, 2016) data from humans. Computational models simulate the PSTH for one or more model ANFs in response to an arbitrary stimulus and therefore offer a rigorous test of the generalization of the Goldstein and Kiang (1958) convolution model for simulating CAPs elicited by a wide variety of acoustic stimuli. The objective of this study is to determine the extent to which this convolution model accounts for human CAPs evoked by a wide range of acoustic stimuli for simulations where *P*(*t*) is defined by PSTHs obtained from two well-established computational models of the human auditory periphery.

## II. GENERAL METHODS

The human CAP was simulated by convolving an estimate of the population post-stimulus time histogram (PPSTH) with a UR (Goldstein and Kiang, 1958):

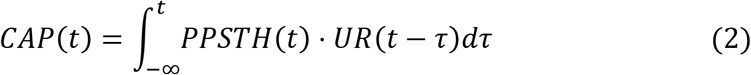

where *t* denotes time and *τ* is an integration variable. The UR represents the waveform of the volume-conducted response resulting from a single-unit action potential, as observed from the site of the recording electrode (Dau, 2003; Ronne *et al.*, 2012). The PPSTH is defined as the across-CF sum of the PSTHs from a population of ANFs (Kiang *et al.*, 1976).

### A. Unit response (UR)

Several UR waveforms have been proposed from animal (Teas *et al.*, 1962; Versnel *et al.*, 1992; McMahon and Patuzzi, 2002; Chertoff, 2004) and human studies (Elberling, 1976). The latter was used in this study given that Elberling (1976) derived this UR from evoked-potential measurements in normal-hearing adult humans, and preliminary simulations revealed that this UR produced CAPs that were qualitatively more similar to empirically recorded human CAPs compared to other candidate URs (e.g., Versnel *et al.*, 1992; Chertoff, 2004). The UR from Elberling (1976) can be accounted for by the following analytic expression:

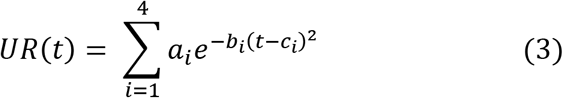

where the coefficients *a_i_, b_i_, c_i_* are given in Table 1. A comparison between this UR and the fit associated with this expression is provided in Figure 1.

**Figure 1:**
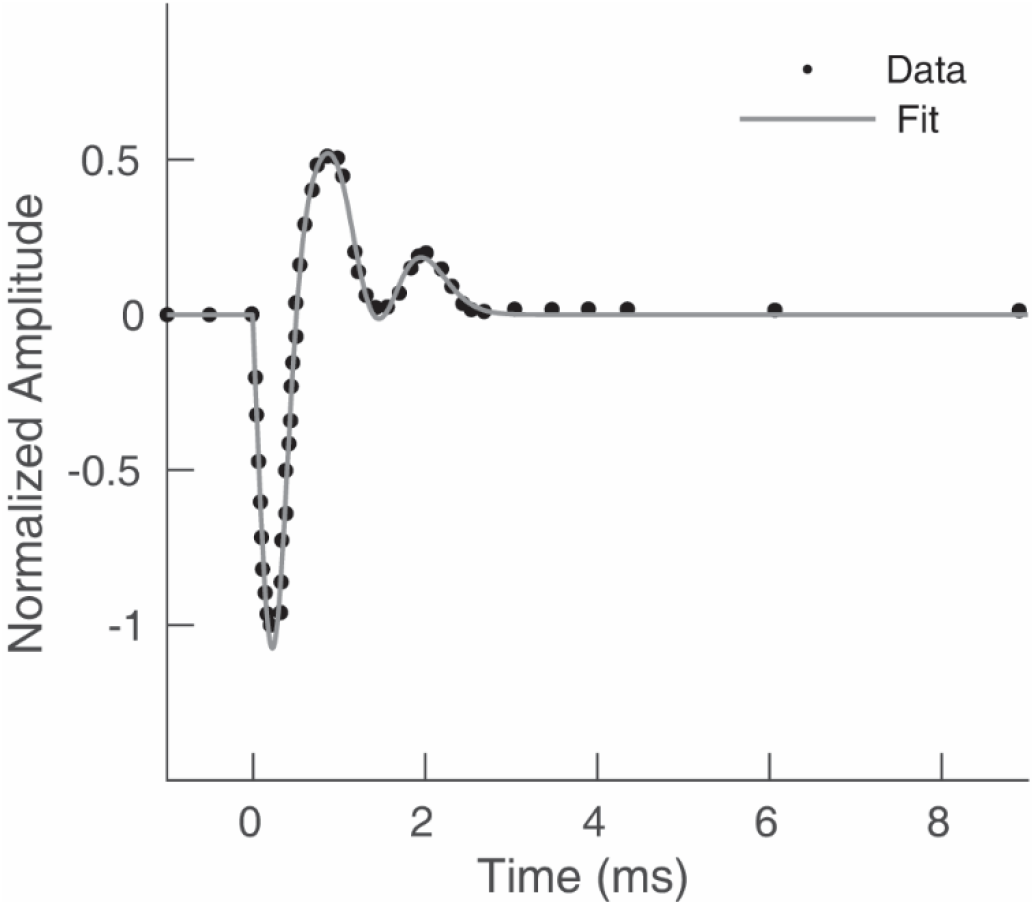
Comparison of the human UR (black dots) estimated by Elberling (1976) and the fit (gray curve) associated with the analytic expression shown in Equation 3.

**Table 1.** The coefficients of the analytic expression (3) for the unitary response.

| Coefficient | Value | Coefficient | Value | Coefficient | Value |
| --- | --- | --- | --- | --- | --- |
| $a_1$ | -49.65299 | $b_1$ | 2.14504 | $c_1$ | 0.83368 |
| $a_2$ | 44.87946 | $b_2$ | 1.96097 | $c_2$ | 0.81179 |
| $a_3$ | 5.51925 | $b_3$ | 4.07791 | $c_3$ | 0.92306 |
| $a_4$ | -2.27615 | $b_4$ | 13.42128 | $c_4$ | 0.19478 |

### B. Population post-stimulus time histogram (PPSTH)

Two well-established computational models of the peripheral auditory system were used to generate the PPSTH from the PSTHs of individual ANFs. The first model was described by Zilany *et al.* (2014) – henceforth referred to as the “Zilany model” – which is a phenomenological model of the auditory periphery that relies on dynamically-controlled nonlinear bandpass filters to account for cochlear processing and incorporates exponential and power-law dynamics to simulate the synapses between the IHCs and ANFs. This model accounts for many physiological properties of the human auditory periphery such as middle-ear filtering (Bruce *et al.*, 2003), and cochlear compression and suppression (Zhang *et al.*, 2001). This model simulates hearing loss due to dysfunctional IHCs and OHCs (e.g., Swaminathan and Heinz, 2011), and can be used to simulate loss of ANFs due to aging and noise exposure (Encina-Llamas *et al.*, 2019). Further, the Zilany *et al.* (2014) model and previous and more recent versions of this model (e.g., Bruce *et al.*, 2018) have been successful at accounting for human psychophysics (e.g., Maxwell *et al.*, 2020) and speech perception (e.g., Haro *et al.*, 2020). The second model used in this study was developed by Verhulst *et al.* (2018a) – henceforth referred to as the “Verhulst model,” – which is a biophysical model that employs a nonlinear transmission line consisting of resonators coupled by cochlear fluids to model cochlear processing. In addition to its successful history of simulating otoacoustic emissions (OAEs), the Verhulst model and its precursor (Verhulst *et al.*, 2015) have been used to simulate a variety of human evoked potentials, such as the auditory brainstem response (ABR) (Verhulst *et al.*, 2015), and the envelope-following response (Verhulst *et al.*, 2018b). Moreover, this model accounts for impaired hearing due to hair cell damage (Verhulst *et al.*, 2016) and cochlear synaptopathy (e.g., Drakopoulos *et al.*, 2022). Given that the Zilany and Verhulst models have been successful at accounting for many aspects of auditory physiology, electrophysiology, and perception in humans, it was hypothesized that these models would not differ substantially in their ability to simulate human CAPs. Thus, the selection of these two models was not intended to provide a rigorous between-model comparison of CAP predictions. Rather, the objective was to test whether the convolution framework of Goldstein and Kiang (1958) combined with a physiologically realistic estimation of ANF activity could account for human CAPs evoked by a wide range of stimuli for at least two established computational auditory models.

To simulate the human PPSTH, the stimuli for a given experiment were generated and presented to 500 CFs spaced logarithmically from 0.25 to 20 kHz (Zilany model), or from 0.025 to 20 kHz (Verhulst model), which are the frequency ranges available for the “humanized” versions of these models. For each CF, fibers with different SRs of discharge were simulated and their responses were weighted according to the ANF distribution of high-, medium, and low-SR categories described by Liberman (1978): 61%, 23%, and 16% for high-, medium-, and low-SR fibers, respectively. Simulated PSTHs were obtained for condensation and rarefaction polarities of each stimulus. This alternating-polarity approach results in cancellation of the CM when responses are averaged across several sweeps (e.g., Henry, 1995). Simulated PSTHs for each model CF were obtained and averaged across 200 repetitions (100 condensation, 100 rarefaction) for the stimuli of each simulated experiment. Finally, the PPSTH was constructed by summing the average PSTHs of high-, medium- and low-SR fibers of the 500 simulated CFs, resulting in a single waveform.

### C. Simulated experiments

The following experiments were simulated and compared to CAPs recorded from human subjects with normal hearing: 1) CAPs evoked by acoustic clicks as a function of click level (Simpson *et al.*, 2020), 2) CAPs evoked by upward frequency chirps as a function of chirp level, 3) narrowband CAPs derived using the high-pass masking technique for CAPs elicited by clicks (Eggermont, 1979b) and TBs (Eggermont *et al.*, 1976), and 4) CAPs evoked by sinusoidally amplitude-modulated (AM) carriers as a function of modulation frequency (Chen and Jennings, 2022). To facilitate comparison with human-subject data, simulated CAP waveforms were scaled to match the baseline-to-peak amplitude of the largest CAP amplitude present in the human-subject dataset associated with each simulated experiment.

## III. SIMULATION I: INTENSITY SERIES FOR AN ACOUSTIC CLICK

### A) Rationale and Methods

An electrical pulse (i.e., click) delivered to an acoustic transducer is one of the most common auditory stimuli used to assess the function of the auditory system. Clicks are broadband transient stimuli that result in synchronous activity from a population of ANFs, thereby eliciting a CAP. When measured across a range of stimulus intensities, the click-evoked CAP provides an estimate of hearing sensitivity for sounds between 2000-4000 Hz (Schoonhoven *et al.*, 1995). The simulations in this section involved predicting CAP waveforms elicited by a 100-*μs* click for intensities ranging from 40 to 110 dB peSPL and comparing these simulations with human-subject data from Simpson *et al.* (2020). In addition to their main experiment, Simpson *et al.* (2020) reported a supplementary experiment where CAPs were measured for 100-*μs* clicks from 40 to 110 dB peSPL in four participants. The human-subject data simulated here included data from these four participants and an additional seven participants – not included in (Simpson *et al.*, 2020) – for whom click-evoked CAPs were measured in the same laboratory. To account for the frequency response of the transducer (ER3C, Etymotic Research, Elk Grove Village, IL) and the resonance of the human outer ear, the click presented to the Zilany and Verhulst models was the average (N=11) waveform measured from the ear canal (ER-7C probe microphone) in response to a 100-μs click presented at 110 dB peSPL. This average waveform was scaled to the desired peSPL (e.g., 70 dB peSPL) before being presented to each model. The simulated CAP waveforms were normalized by the N_1_ amplitude of the largest (1.3 μV) CAP observed in the human-subject dataset, which was for the 110 dB peSPL click. Specifically, the un-normalized model CAP waveform associated with click intensity X was multiplied by 1.3 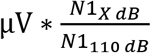 where N1_XdB_ and N1_110dB_ are the N_1_ amplitudes of the un-normalized model CAP waveforms for X dB peSPL and 110 dB peSPL clicks, respectively.

### B) Results and discussion

Fig. 2 shows the average click-evoked CAP waveforms recorded from 11 normal-hearing human subjects (Fig. 2A), and simulated waveforms based on the Zilany (Fig. 2B) and Verhulst (Fig. 2C) models. The human CAP evoked by a high-level click is characterized by a negative deflection (N_1_) followed by two positive deflections (P_1_ and P_2_) with an intermediate smaller negative deflection (N_2_). As the intensity of the click decreases, the N_1_ amplitude decreases, the N_1_ latency increases, and a broader biphasic response is observed. Simulated CAP waveforms are similar to those for human subjects but show some inconsistencies including the observations that the SP is not present for both models, and the morphology of the predicted P_2_ and N_2_ potentials is broader and less resolved than the human-subject data. The SP is observed in the human-subject waveforms as a small shoulder or bump occurring just before N_1_. These deviations from the human data can be explained by the fact that the primary generators of the SP are the cochlear hair cells (Durrant *et al.*, 1998), which were not simulated here. Moreover, the P_2_ and N_2_ potentials are thought to originate from activity of the cochlear nucleus or a portion of the auditory nerve located remotely from the spiral ganglion neurons (Brown and Patuzzi, 2010). Such generators (e.g., cochlear nucleus) were not simulated for the current Zilany and Verhulst model predictions.

**Figure 2:**
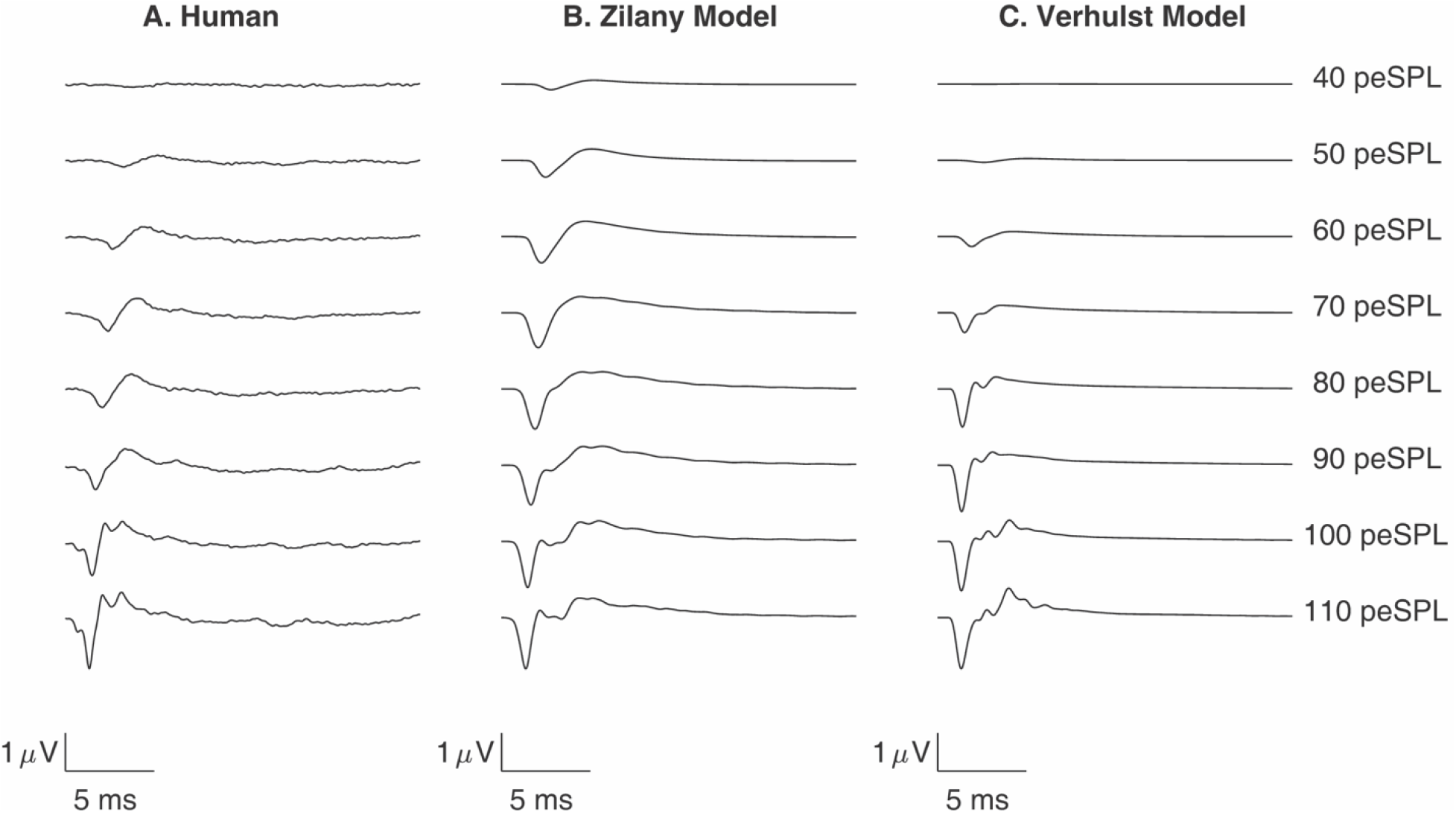
CAPs evoked by 100-μs acoustic clicks presented at intensities from 40 to 110 dB peSPL for human (A) and simulated waveforms from the Zilany (B) and Verhulst (C) models.

Human-subject N_1_ amplitudes and latencies are quantified in Fig. 3A, D. N_1_ baseline-to-peak amplitudes reach a maximum of 1.3 μV, grow rapidly between 40-70 dB and 90-110 dB peSPL, and more slowly for intermediate levels (Schoonhoven *et al.*, 1995). N_1_ latencies span a range of 2.75 ms and are characterized by a decrease in slope with increasing click level (Fig. 3D). Elberling (1974) derived narrowband responses to interpret the level-dependent effects of N_1_ amplitude and latency for the click-evoked CAP. He concluded that CAPs evoked by a lower-level click (75 dB peSPL) were produced *“mainly by components originating more apically in the cochlea (2-4 kHz area),”* while CAPs evoked by a higher-level click (95 dB peSPL) were produced *“mainly by components originating in the basal part of the cochlea (2-4 kHz area).”* Thus, the level-dependent changes in N_1_ amplitude and latency are consistent with the cochlear response spreading basalward with increasing click level (i.e., upward spread of excitation), as concluded by Elberling (1974) and other studies of the CAP (Eggermont *et al.*, 1976; Schoonhoven *et al.*, 1995).

**Figure 3:**
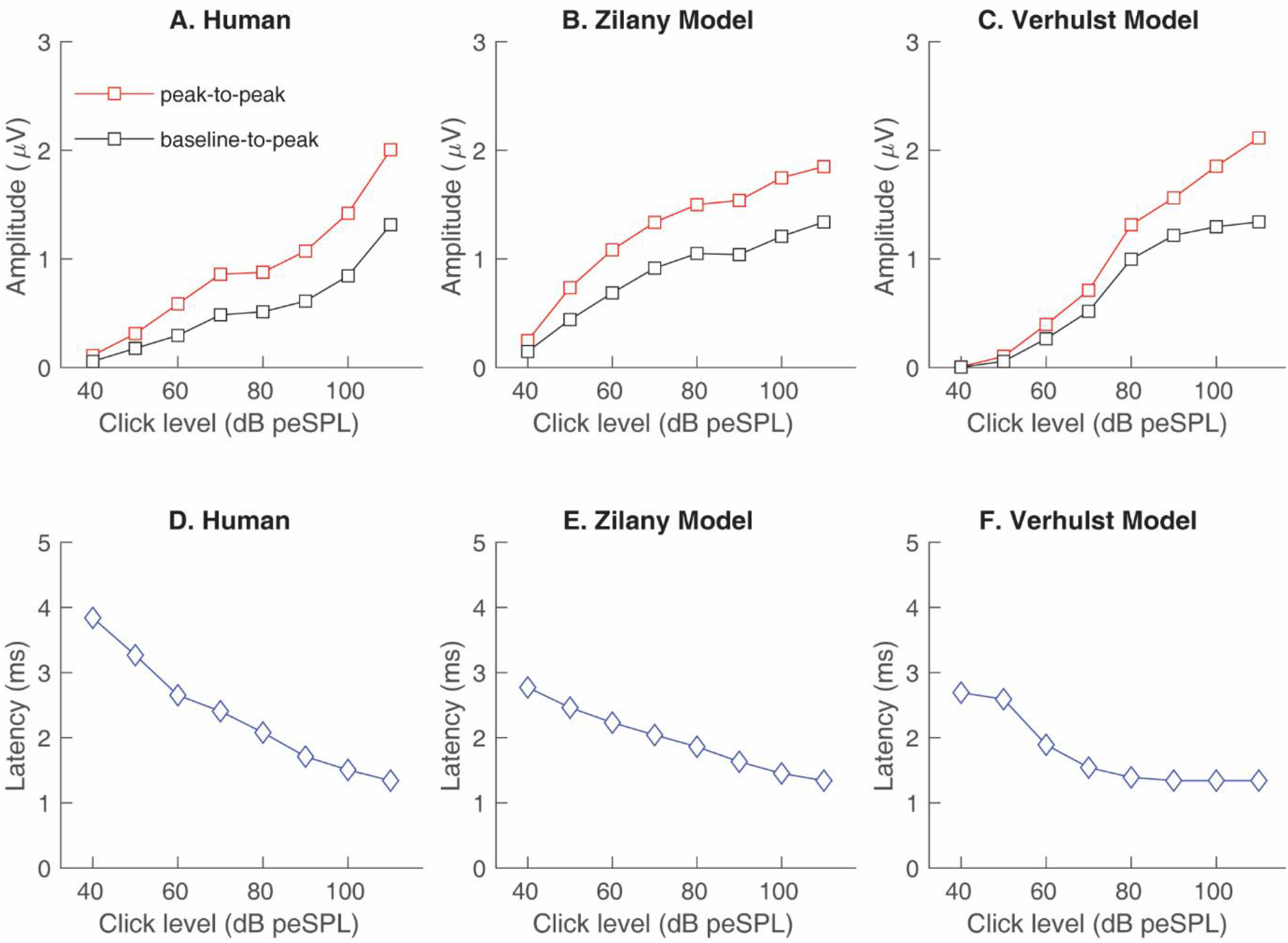
N_1_ amplitude and latency as a function of click intensity. A-C: N_1_ peak-to-peak (red curves) and baseline-to-peak (black curves) amplitudes for the average human CAPs (A), and simulated CAPs associated with the Zilany (B) and Verhulst (C) models. D-F: N_1_ latency for the average human CAPs (D), and simulated CAPs associated with the Zilany (E), and Verhulst (F) models.

The model simulations account for human CAP amplitude and latency trends as quantified in Fig. 3 B, C, E, F, which shows N_1_ peak-to-peak and baseline-to-peak amplitudes (Fig. 3, top row), and latencies (Fig. 3 bottom row) as a function of click intensity. Specifically, simulated CAPs obtained with the Zilany and Verhulst models show an increase in N_1_ amplitude and a decrease in N_1_ latency as the click level increases. Compared to the human subject click-evoked CAPs, the predicted CAPs grow faster in amplitude between 40-80 dB peSPL and slower in amplitude between 90-110 dB peSPL. Furthermore, the range of latencies is smaller for simulated CAPs compared to human-subject CAPs. These differences between simulated and human-subject CAPs for clicks may be related to the ability of the models to account for the details of the upward spread of excitation of the human cochlea, as considered in the Discussion section.

## IV. SIMULATION II: INTENSITY SERIES FOR AN ACOUSTIC CHIRP

### A) Rationale and Methods

A rising frequency chirp is thought to result in greater across-CF synchrony of ANFs compared to a click by compensating for the basilar membrane traveling wave delay (Shore and Nuttall, 1985). Given this advantage over clicks, chirps are often used for eliciting the ABR in clinical practice (e.g., Bargen, 2015). Furthermore, chirps may be a useful stimulus for identifying cochlear synaptopathy (Earl, 2015). Chertoff *et al.* (2010) compared CAPs elicited by a 100-μs click with those elicited by a rising frequency chirp in normal-hearing humans, where the chirp had the same spectrum as the click and the instantaneous frequency of the chirp was based on traveling wave estimates from derived narrowband CAPs measured in human subjects (Eggermont, 1979a). The instantaneous frequency of the chirp used by Chertoff *et al.* (2010) is level independent and contrasts with level-dependent chirps designed to optimize Wave-V amplitude of the ABR across a wide range of levels (Elberling and Don, 2010). Chirp-evoked CAPs, using the chirp from Chertoff *et al.* (2010), were simulated with the Zilany and Verhulst models and compared with empirical CAPs evoked by clicks and chirps from 12 young adults with normal hearing (<25 dB HL 250-8000 Hz) for intensities of 50, 60, 70, 85, and 100 dB peSPL. These data were obtained as part of an ongoing study in the Auditory Perception and Physiology Laboratory at the University of Utah and followed the procedures described by Simpson *et al.* (2020) for measuring ECochG with a custom TM electrode. The simulated CAPs included chirp intensities ranging from 40 to 110 dB peSPL. The click and chirp waveforms delivered to the models were those measured (System 824, Larson-Davis) at the output of a 2-cc coupler (AEC203 Larson-Davis) when presented through ER3C earphones, which were the earphones used for the human-subject experiment. The simulated CAP waveforms were normalized by the average N_1_ amplitude (1.0 μV) observed in human subjects for a 100 dB peSPL click. Specifically, the un-normalized model CAP waveform associated with click (or chirp) intensity X was multiplied by 1.0 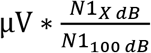, where N1_XdB_ and N1_100dB_ are the N_1_ amplitudes of the un-normalized model CAP waveforms for an X dB peSPL click or chirp and the 100 dB peSPL click, respectively.

### B) Results and discussion

Fig. 4A shows the average click- and chirp-evoked CAP waveforms (gray and black lines, respectively) recorded from 12 normal-hearing human subjects. The major finding is that the N_1_ amplitude is greater for chirp-than for click-evoked CAPs (i.e., chirp benefit) for the lowest levels (50 and 60 dB peSPL), consistent with a similar chirp benefit reported by Chertoff *et al.* (2010) and Smith *et al.* (2017). For the highest level tested (100 dB peSPL), the opposite effect is observed, where average N_1_ amplitudes were larger for the click-than the chirp-evoked CAP. A similar lack of a chirp benefit for high levels of stimulation was reported for Wave-I of the ABR when using a level-independent chirp comparable to the one used here (“CE-chirp,” Morimoto *et al.*, 2019). Differences in N_1_ latency for the human click- and chirp-evoked CAPs can be observed in Fig. 4A, where the time corresponding to 0 ms is the *offset* of the stimulus. Specifically, the chirp-evoked N_1_ appears earlier in time than the click-evoked N_1_ for all but the lowest stimulus level (50 dB peSPL).

**Figure 4:**
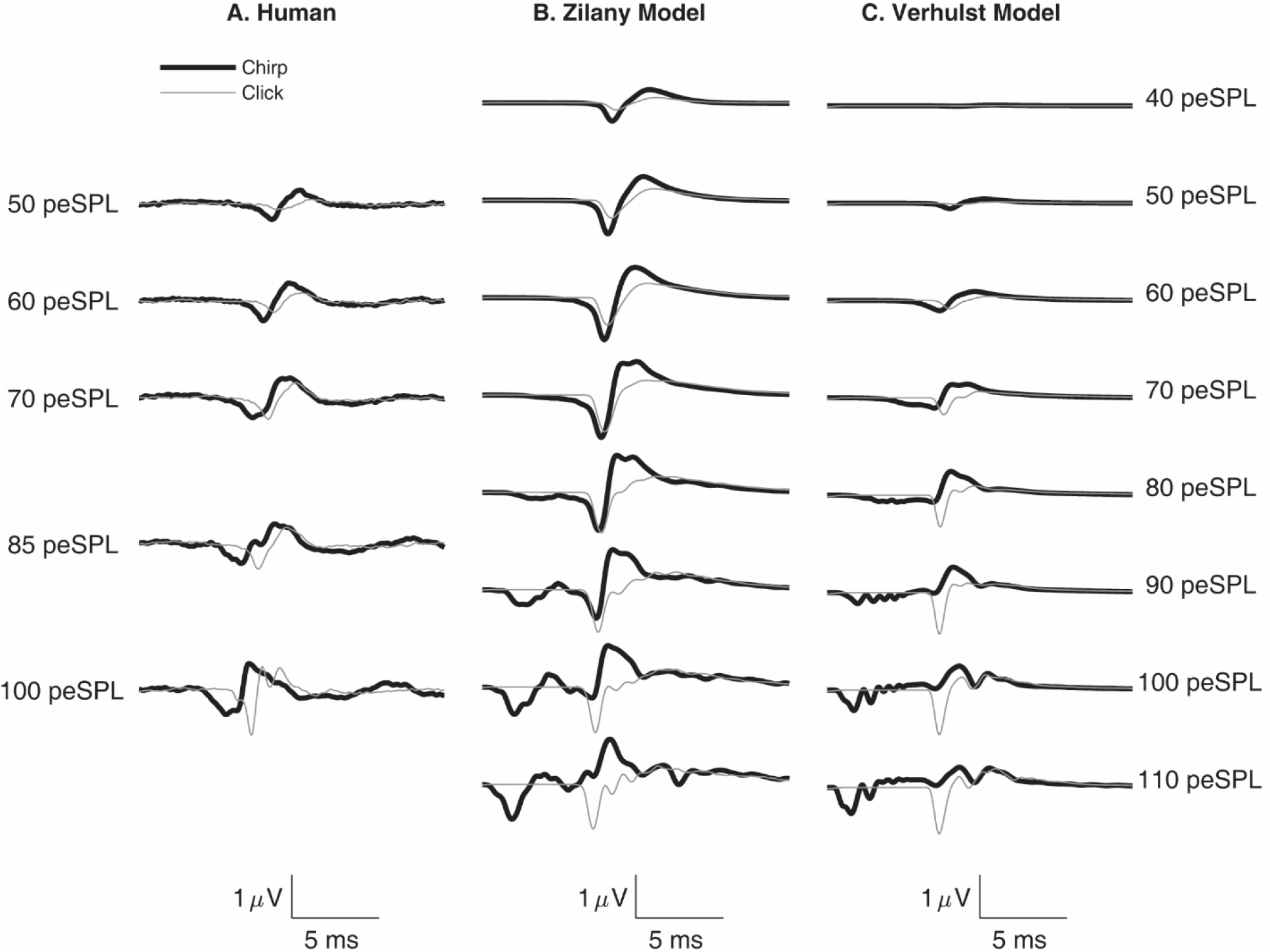
Human and simulated CAPs evoked by a rising-frequency chirp (black curves) or click (gray curves) as a function of stimulus intensity. A: the average chirp- or click-evoked CAPs from 12 normal-hearing subjects. B: The simulated chirp- and click-evoked CAPs associated with the Zilany model. C) The simulated chirp- and click-evoked CAPs associated with the Verhulst model.

Fig. 4B and Fig. 4C display simulated click- and chirp-evoked CAPs obtained with the Zilany and Verhulst models. CAPs simulated from both models exhibit a chirp benefit for lower levels (Zilany, < 80 dB peSPL; Verhulst, <70 dB peSPL), and the absence of this benefit for higher levels for baseline-to-peak N_1_ amplitudes. The prediction of a chirp benefit for lower-level chirps is consistent with the human-subject CAPs; however, simulations predict that this benefit will be larger or smaller than that observed in human subjects for the Zilany and Verhulst models, respectively. Similarly, consistent with the human-subject CAPs, simulations predict a broadening of the chirp-evoked CAP with increasing level; however, these simulations incorrectly predict the emergence of an early-latency negative potential when the level is greater than ~80 dB peSPL. Finally, compared to the human-subject CAPs, the amplitude of N_1_ decreases at a slower (Zilany model) and faster (Verhulst model) rate with decreasing chirp intensity. This difference in rate suggests that predicted N_1_ thresholds will be lower or higher than the human-subject CAPs for the Zilany and Verhulst models, respectively.

The top row of Fig. 5 displays the N_1_ amplitudes associated with the empirical and simulated CAPs from Fig. 4. The human CAP exhibits a nearly constant N_1_ amplitude as a function of chirp intensity (Fig. 5A). Conversely, simulations incorrectly predict non-monotonic growth of N_1_ amplitudes (Fig. 5B, C), where amplitudes increase, decrease, and increase again as a function of chirp level. Although this non-monotonic growth of N_1_ amplitude is absent for the human-subject CAPs, previous studies have reported an increase then decrease in ABR Wave-V amplitudes as a function of chirp level when measurements extended to ~115 dB peSPL (Kristensen and Elberling, 2012). The human-subject data presented here was limited to a maximum level of 100 dB peSPL. This suggests that differences in the maximum level (i.e., 100 dB peSPL; 115 dB peSPL) may explain why nonmonotonic amplitude growth for chirp-evoked responses from human subjects was not observed in this study compared to previous ABR studies.

**Figure 5:**
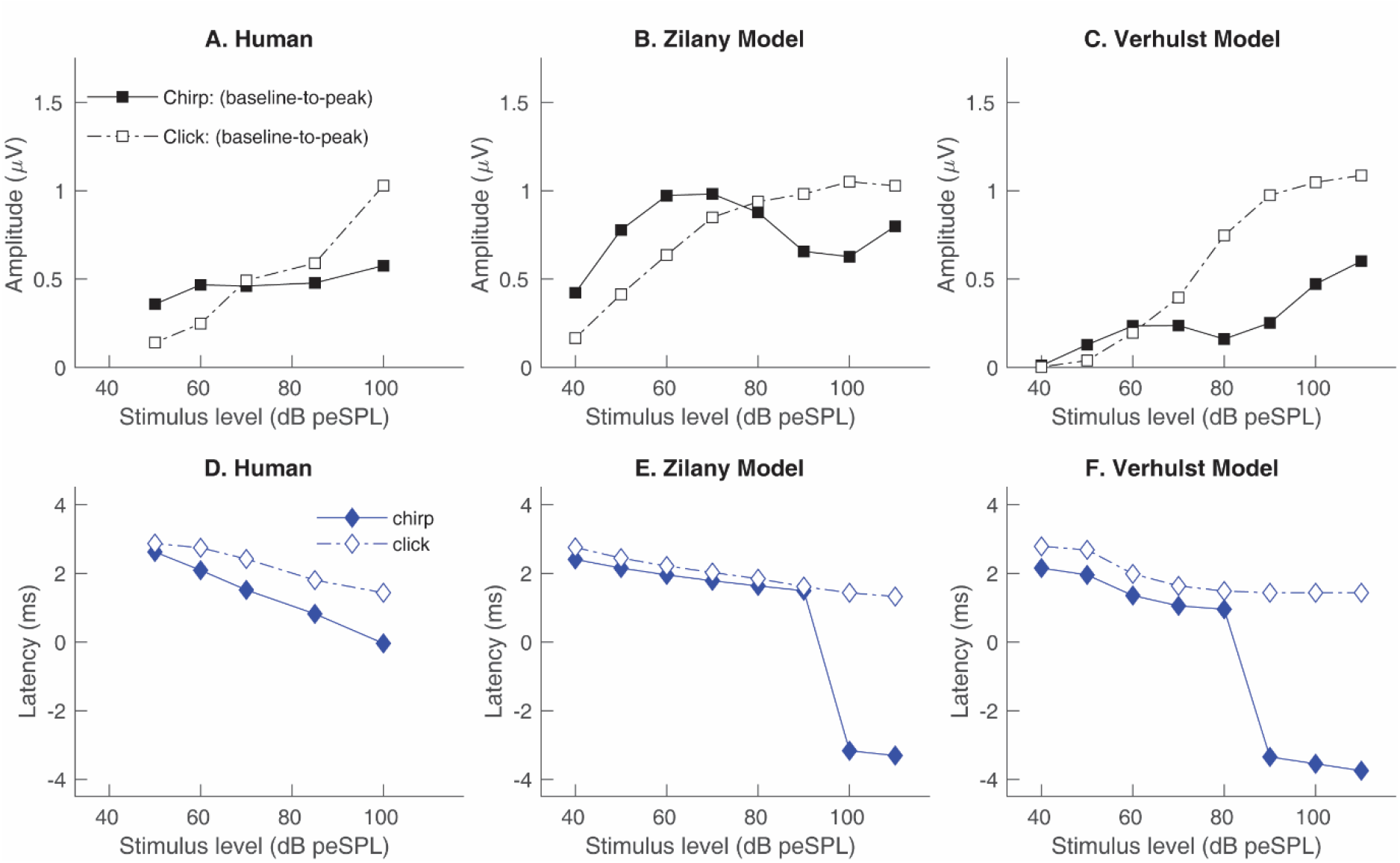
N_1_ amplitude and latency for human and simulated CAPs for chirps (solid curves) and clicks (dashed curves) as a function of stimulus intensity. A-C: baseline-to-peak N_1_ amplitudes for the average human CAPs evoked by clicks and chirps (A), simulated CAPs associated with the Zilany model (B), and the Verhulst model (C). D-F: N_1_ latency with respect to the offset of the stimulus for the average human CAPs (D), or simulated CAPs associated with the Zilany (E) or Verhulst (F) models.

The bottom row of Fig. 5 displays N_1_ latencies as a function of stimulus intensity for the empirical and simulated CAPs evoked by clicks and chirps. For human subjects, the N_1_ latency-intensity function for chirps is steeper than that for clicks and spans a wider range of latencies for the levels (i.e., 50 – 100 dB peSPL) tested in this study. These characteristics (steeper slope, wider range) are observed for the simulated chirp-evoked N_1_ latencies; however, the simulations incorrectly predict an abrupt decrease in latency around 80 (Verhulst model) or 90 (Zilany model) dB peSPL. This abrupt decrease in latency occurs because the identified N_1_ jumps to an earlier negative potential in the waveform (Fig. 4B, C).

Fig. 6 displays the contribution of model ANFs of differing SRs (black: high SR [HSR], red: medium SR [MSR], and cyan: low SR [LSR]) to the simulated CAP elicited by 60 dB peSPL clicks and chirps for the Zilany (Fig. 6 A, B) and Verhulst (Fig. 6 C, D) models. These figures are inspired by Antoli-Candela and Kiang (1978), who recorded single-unit and CAP responses simultaneously from the cat AN (see their Figs. 11–12). The thick solid lines at the top of each panel display the simulated CAP and PPSTH. Simulations for the click (left column) exhibit increasing delay of the onset response of model ANFs as CF decreases. This delay highlights the ability of these models to account for the finding that traveling wave velocity decreases with decreasing CF (Zerlin, 1969), thereby resulting in greater synchrony of basal compared to apical ANFs (Neely *et al.*, 2003). Simulations for the chirp exhibit an onset response from model ANFs that is more temporally aligned (right column) across CFs than that for the click. This alignment was observed for the responses of both models and is consistent with the hypothesis that a chirp compensates for the temporal smearing of the neural activity that is associated with the cochlear traveling wave delay (Dau *et al.*, 2000). Both models predict a chirp benefit for this level (60 dB peSPL), as shown by the simulated CAPs.

**Figure 6:**
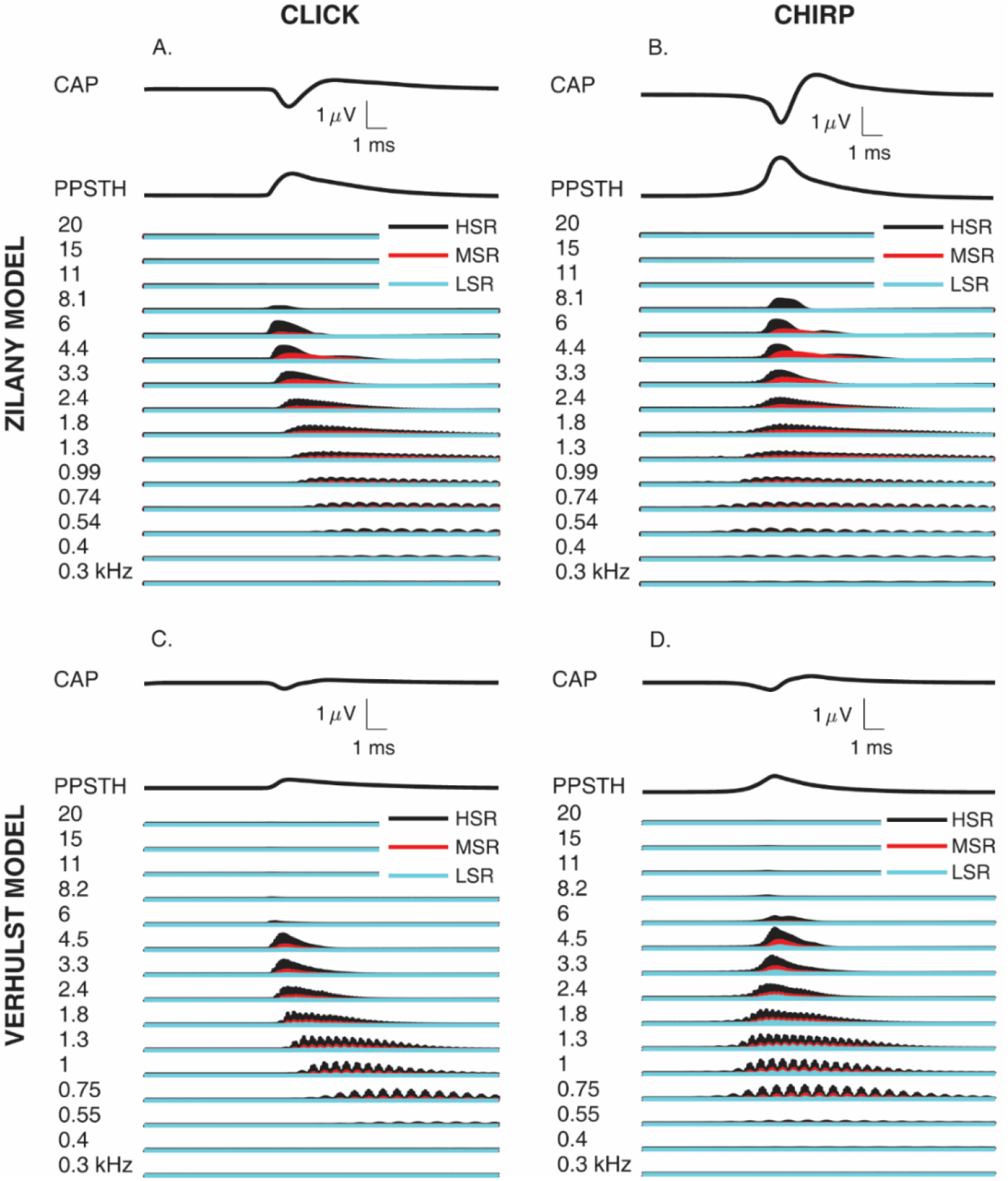
Analysis of single-unit contributions to the simulated 60 dB peSPL click- (left panel) and chirp-evoked (right panel) CAPs using Zilany (top row panels) and Verhulst (bottom row panels) models. For each panel, the predicted CAP and PPSTH waveforms are the top and middle traces, respectively. The lower set of traces are the predicted PSTHs for ANFs with CFs indicated on the left margin. Black, red, and cyan traces are associated with HSR, MSR, and LSR ANFs, respectively.

**Figure 7:**
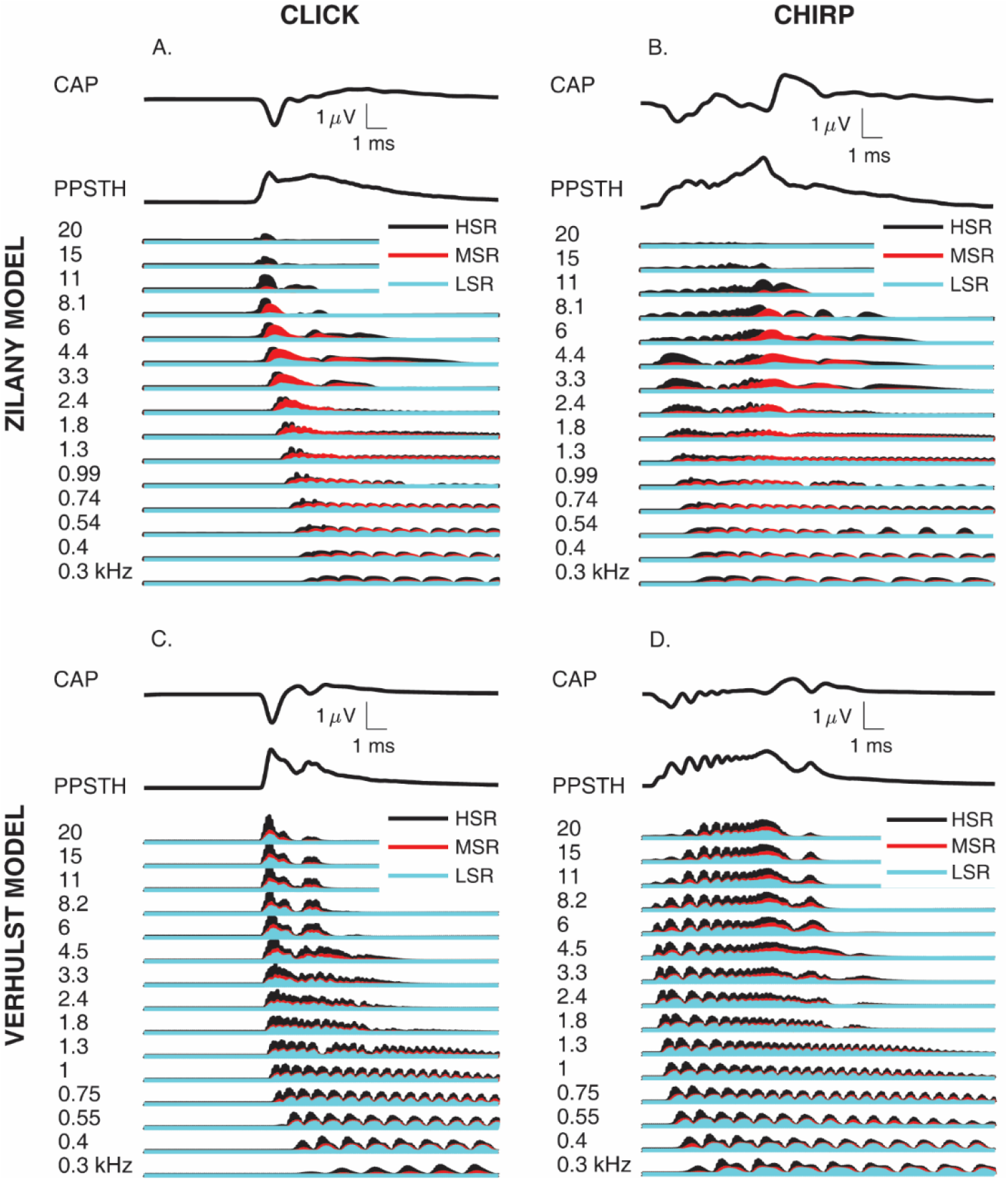
Analysis of single-unit contributions to the simulated 100 dB peSPL click- (left panel) and chirp-evoked (right panel) CAPs using Zilany (top row panels) and Verhulst (bottom row panels) models. For each panel, the predicted CAP and PPSTH waveforms are the top and middle traces, respectively. The lower set of traces are the predicted PSTHs for ANFs with CFs indicated on the left margin. Black, red, and cyan traces are associated with HSR, MSR, and LSR ANFs, respectively.

**Figure 8:**
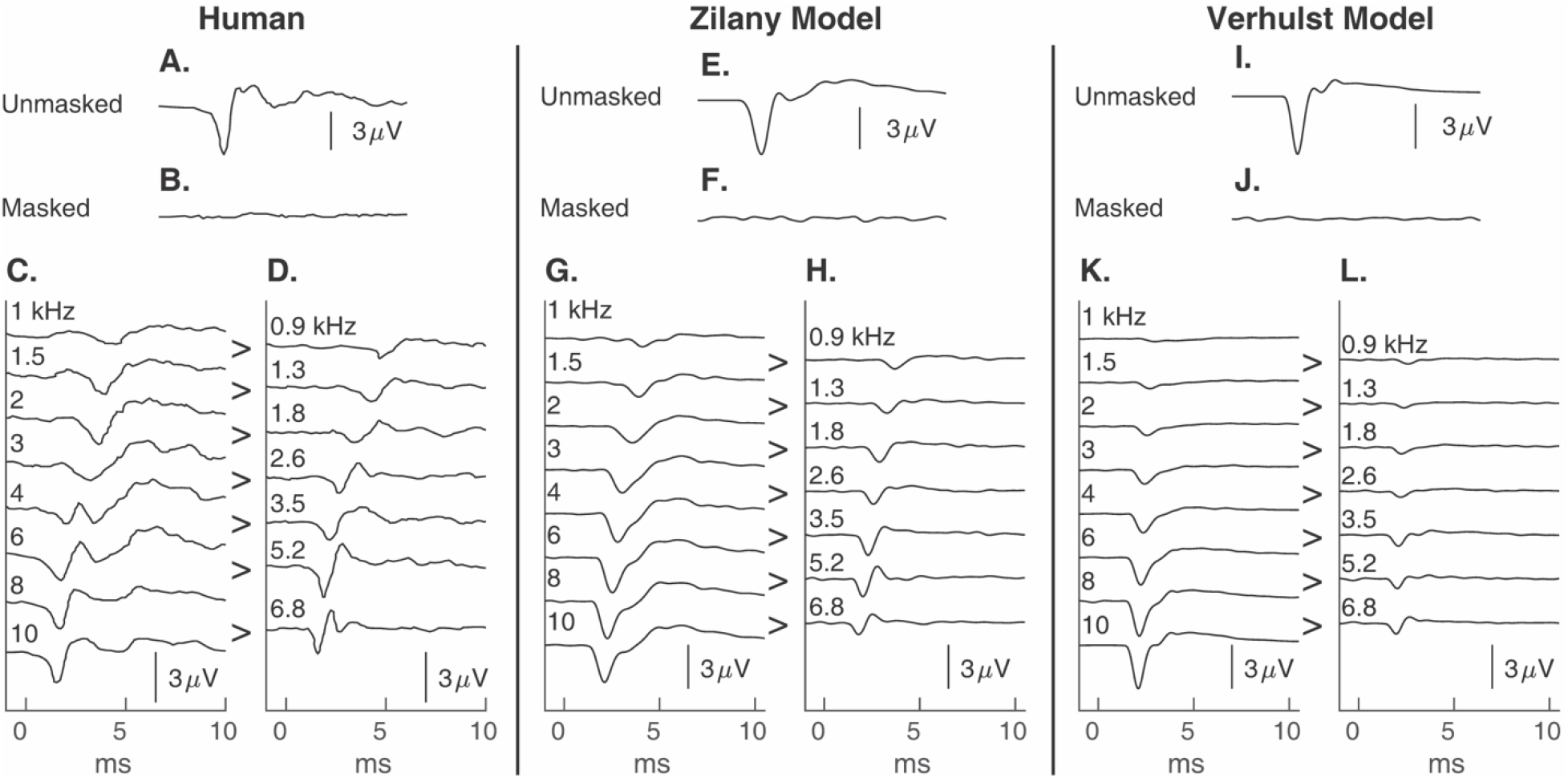
Human (A-D) (adapted from Eggermont, 1979b) and simulated derived narrowband CAPs elicited by a 90 dB peSPL acoustic click obtain with the Zilany (E-H) Verhulst (I-H) models. A,E,I: unmasked CAPs. B,F,J: fully masked CAPs. C,G,K: high-pass masked CAPs shown with the corresponding cutoff frequency. D,H,L: derived narrowband CAPs with the corresponding central frequency.

**Figure 9:**
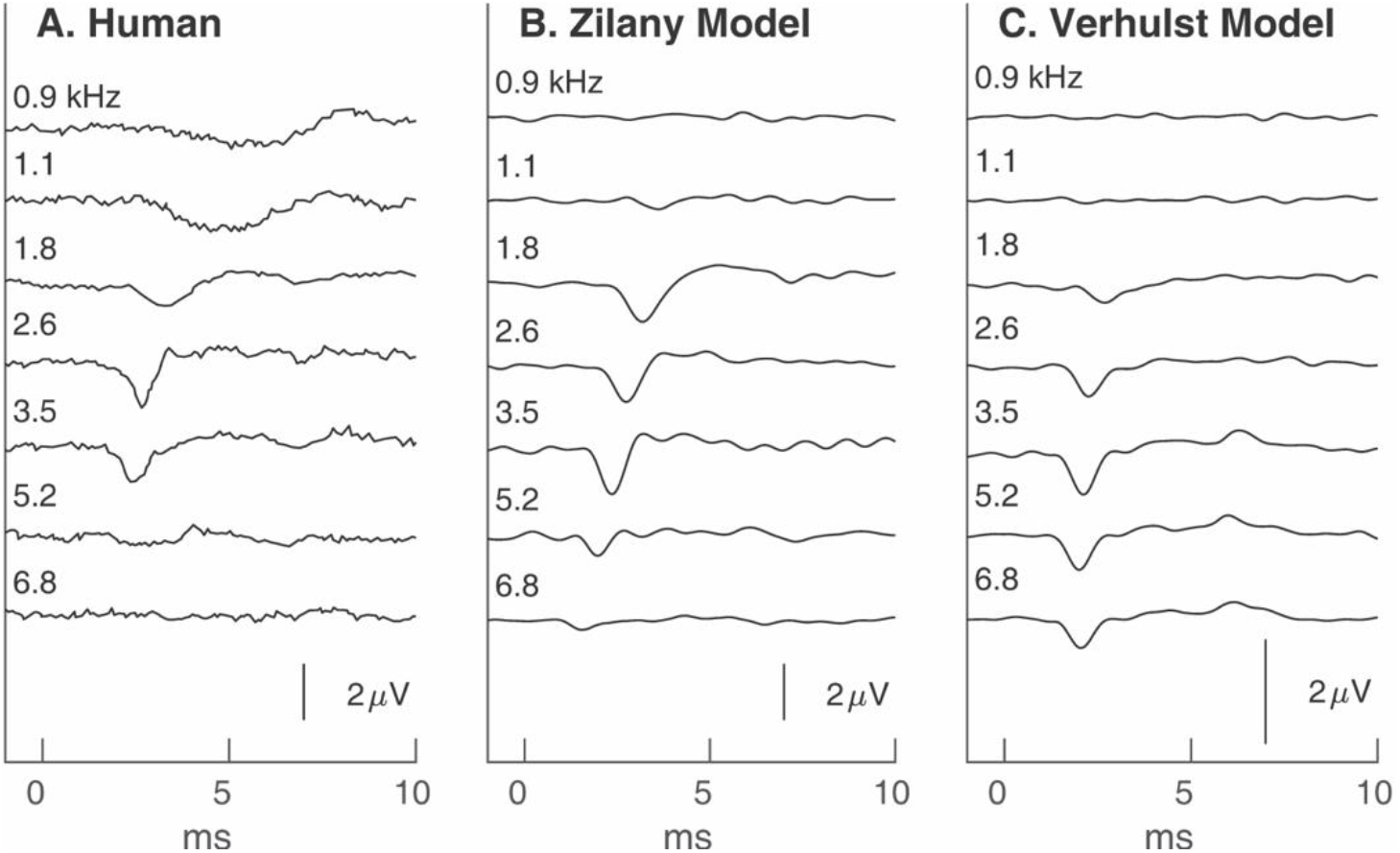
Human (adapted from Eggermont, 1976) and simulated derived narrowband CAPs elicited by a 75-dB peSPL, 2000-Hz TB. The human (A) and simulated derived narrowband CAPs obtained using the Zilany (B) and Verhulst (C) models are shown with the corresponding central frequency.

**Figure 10:**
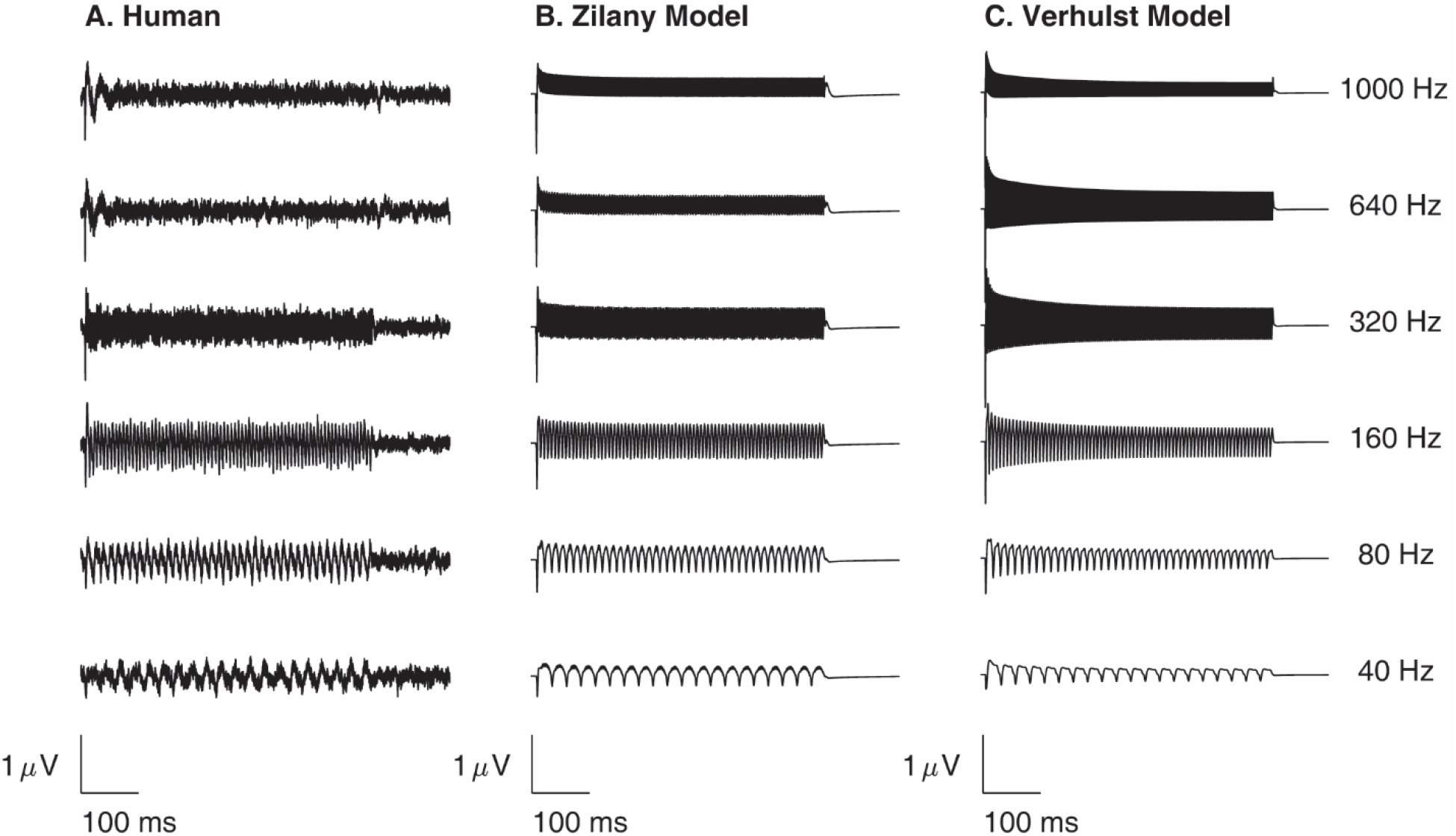
Human and simulated CAP_ENV_ evoked by a 3000-Hz, AM carrier for modulation frequencies ranging from 40 to 1000 Hz. A) Average human CAP_ENV_ from eight normal-hearing subjects (Chen and Jennings, 2022). B) Simulated CAP_ENV_ associated with the Zilany model. C) Simulated CAP_ENV_ associated with the Verhulst model.

**Figure 11:**
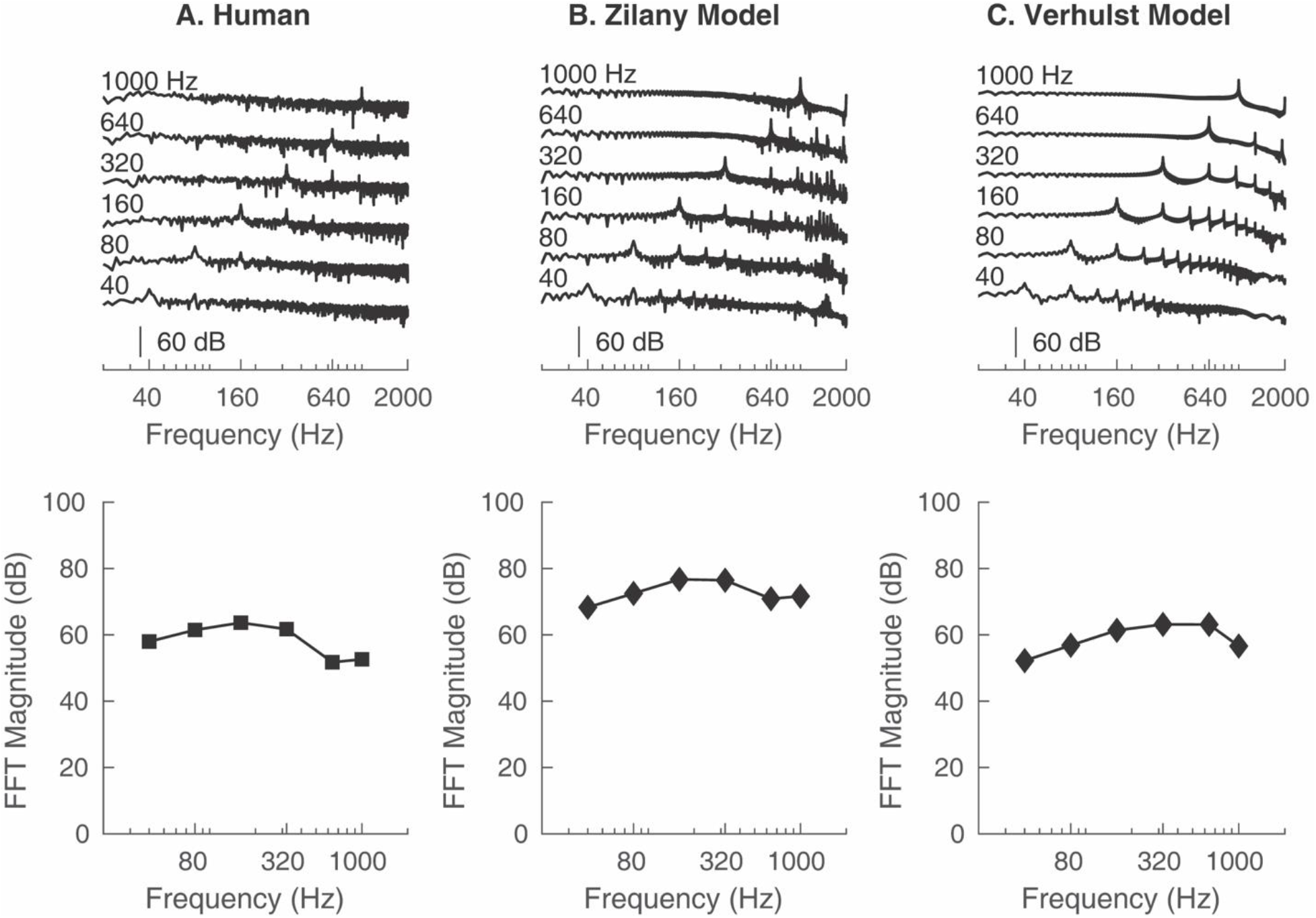
The spectra of the human (A) and simulated (B, C) CAP_ENV_ (top row) and the magnitudes of the associated modulation frequencies (bottom row).

**Figure 12:**
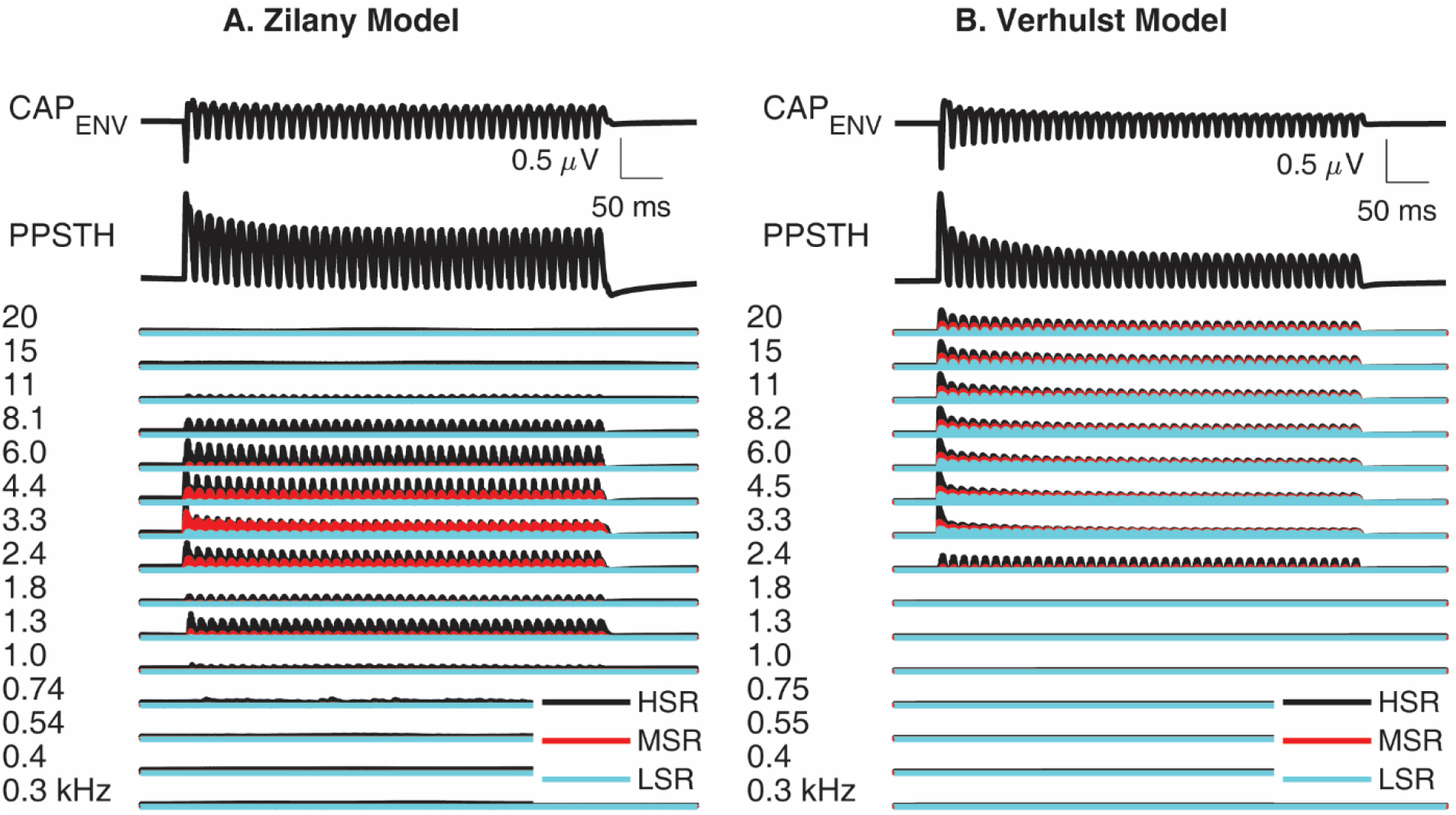
Analysis of single-unit contributions to CAP_ENV_ for simulations associated with the Zilany (A) and Verhulst (B) models. For each panel, the predicted CAP_ENV_ and PPSTH waveforms are the top and middle traces, respectively. The lower set of traces are the predicted PSTHs for ANFs with CFs indicated on the left margin. Black, red, and cyan traces are associated with HSR, MSR, and LSR ANFs, respectively.

Fig. 7 displays the contribution of model ANFs to the simulated CAPs elicited by 100 dB peSPL clicks and chirps and follows the same format as Fig. 6. Similar to the 60 dB peSPL click, ANF responses to the 100 dB peSPL click exhibit a delay in onset activity, particularly for CFs below 1.8 kHz. Unlike the 60 dB peSPL chirp, however, the 100 dB peSPL chirp was ineffective at aligning the responses of ANFs across CFs, resulting in smaller N_1_ amplitudes for chirp-compared to click-evoked CAPs. The absence of a chirp benefit for high stimulus levels was addressed for the ABR by Kristensen and Elberling (2012), who hypothesized that upward spread of excitation temporally smears the responses of ANFs by exciting high-CF fibers by both low- and high-frequency components of the chirp. Model simulations support this hypothesis as nearly all model ANFs are stimulated throughout the duration of the chirp. This contrasts with simulations for 60 dB peSPL (Fig. 6), where ANF activity was limited to near the chirp’s offset. Another consequence of upward spread of excitation is the adaptation of ANFs. Specifically, high-CF model fibers were stimulated during the upward frequency chirp for a wide range of frequencies below these fibers’ CFs. Stimulation by frequencies lower than a given CF adapted the fiber and resulted in a diminished firing rate when the instantaneous frequency of the chirp equaled the fiber’s CF. This phenomenon can be illustrated by comparing peak firing rates in response to 100 dB peSPL clicks and chirps for ANFs with CFs >1.3 kHz (e.g., Fig. 7, lines labeled HSR).

## V: SIMULATION III: HIGH-PASS MASKING AND DERIVED NARROWBAND CAPS

### A) Rationale and Methods

CAPs elicited by a broadband click or chirp include responses from a population of ANFs with a wide range of CFs. However, CAPs elicited by these broadband stimuli do not reveal how ANFs tuned to a narrow frequency range (e.g., 1700-2200 Hz) contribute to the CAP. High-pass masking techniques have been developed to derive the CAP elicited from ANFs within a narrow range of CFs (i.e., derived narrowband CAPs). Applications for derived narrowband responses include estimating the cochlear traveling wave (Eggermont, 1976; 1979a), evaluating spread of cochlear excitation (Eggermont, 1976; 1979a), and detecting the presence of an eighth-nerve tumor from the “stacked” ABR (Don *et al.*, 2005). Simulations of derived narrowband CAPs were obtained using the high-pass noise technique for a 90 dB peSPL click and a 75 dB peSPL, 2000-Hz TB. These simulations were compared with human-subject data from Eggermont (1979b) and Eggermont (1976) for derived narrowband CAPs evoked by click and TB stimuli, respectively. Simulations were obtained by first determining the spectrum level of a broadband (0 – 10,000 Hz) noise that fully masked the simulated click-evoked CAP. All noises and click stimuli were lowpass filtered with a cutoff-frequency of 10,000 Hz to account for the roll off of the transducer reported by Eggermont (1976). Next, CAPs evoked by clicks were obtained in the presence of high-pass noise with the following cutoff frequencies: 1.0, 1.5, 2.0, 3.0, 4.0, 6.0, 8.0, and 10.0 kHz. The spectrum level of the high-pass noise was maintained at the spectrum level of the broadband noise that fully masked the CAP (Teas *et al.*, 1962). Narrowband CAPs were derived by subtracting waveforms from simulations having adjacent high-pass noise cutoff frequencies (e.g., subtracting the waveforms associated with 3.0 and 2.0 kHz cutoff frequencies derived the narrowband CAP centered on 1.8 kHz). A similar process was used for obtaining derived narrowband CAPs for the 2000-Hz TB, except the 1.4 kHz cutoff frequency was increased to 1.5 kHz, consistent with Eggermont (1976). This subtraction procedure resulted in seven narrowband responses corresponding to the following central frequencies: 0.9, 1.3, 1.8, 2.6, 3.5, 5.2, 6.8 kHz (for clicks), and 0.9, 1.1, 1.8, 2.6, 3.5, 5.2, 6.8 kHz (for TBs). The waveforms for simulated click-evoked CAPs in broadband and high-pass noise were normalized by the N_1_ amplitude of the unmasked click-evoked CAP from Eggermont (1979b), which was 5 μV. A similar normalization was applied to the simulated TB-evoked CAPs, where the reference voltage was 3 μV (Eggermont, 1976). Specifically, the un-normalized click-evoked CAP waveform for a given noise condition was multiplied by 5.0 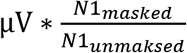 (clicks), or 3.0 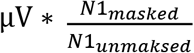 (TB), where N1_masked_ and N1_unmasked_ are the N_1_ amplitudes of the un-normalized model CAP waveforms for a given noise condition (e.g., broadband noise, or high-pass noise) and the unmasked condition, respectively.

### B) Results and discussion

Fig. 8 illustrates the human (Eggermont, 1979b) [A-D] and simulated (Zilany model [E-H], Verhulst model [I-L]) derived narrowband CAPs for a 90 dB peSPL click. CAPs in the unmasked (Fig. 8, A, E, I), fully masked (Fig. 8 B, F, J), and high-pass noise (Fig. 8 C, G, K) conditions are displayed. For CAPs evoked by a click in high-pass noise (Fig. 8 C, G, K), the N_1_ amplitude increases and the latency decreases as the noise cutoff frequency increases. Although the simulations capture these general trends for amplitude and latency, the latency shifts for simulated CAPs are smaller than the human-subject data, particularly for the Verhulst model. Furthermore, simulations do not account for the latency difference between N_1_ and the following broad negative potential (N_2_) for the unmasked CAP (cf. Fig. 8 E, I and Fig. 8 A). This latency difference resulted in the emergence of a double-peaked CAP waveforms for human subjects for noise cutoff frequencies of 4 and 6 kHz (Fig. 8 C). Eggermont (1976) interpreted the N_2_ potential elicited by a 90 dB peSPL click in the unmasked condition as evidence for the contribution of apical ANFs to the CAP waveform. Model simulations predict a similar apical contribution; however, the latency of this contribution is earlier than data from human subjects.

The derived narrowband CAPs for clicks are presented in Fig. 8 D, H, K for waveforms from human-subjects and model simulations. Similar to the results from human subjects (Fig. 8D), derived narrowband CAPs simulated with the Zilany model (Fig 8 H) show a maximum peak-to-peak amplitude at a central frequency of 5.2 kHz. In contrast, the derived narrowband CAPs simulated with the Verhulst model (Fig. 8 L) exhibit monotonically increasing amplitudes with increasing central frequency. For both models, the amplitudes of the derived narrowband CAPs are smaller than those from human subjects.

The monotonic increase in the amplitude of the derived CAPs obtained with the Verhulst model is an expected consequence of subtracting the model waveforms with adjacent high-pass noise cutoff frequencies (Fig. 8 K). These waveforms have similar morphologies and latencies, thus subtracting each pair results in appreciable destructive interference and leaves only the uncanceled component of the waveform that has the greater amplitude.

Figure 9 displays derived narrowband CAPs for the 2000 Hz TB for human subjects [A] (Eggermont, 1976), and model simulations (Zilany model [B], Verhulst model [C]). The largest amplitudes for the derived narrowband CAPs evoked by a TB are near the TB frequency (2000 Hz) for data from human subjects. Specifically, amplitudes are largest for the 1.8, 2.6, and 3.5 kHz bands, and peak at the 2.6 kHz band. A similar result is observed for simulations from the Zilany model; however, these derived narrowband CAPs for these bands have similar amplitudes rather than a peak at 2.6 kHz, as was observed for human-subject data. For simulations from the Verhulst model, the largest amplitudes of the derived narrowband CAPs were observed for the bands ranging from 1.8 to 6.8 kHz. The prediction of an N_1_ for derived narrowband CAPs with central frequencies of 5.2 and 6.8 kHz is inconsistent with the results from human subjects. Additionally, like the narrowband analysis of click-evoked CAPs, the derived narrowband CAPs obtained for a TB with the Verhulst model show an overall smaller magnitude than those for the human-subject data and Zilany simulations.

## VI: SIMULATION IV: AMPLITUDE-MODULATED SINUSOIDAL CARRIERS

### A) Rationale and Methods

The auditory steady-state response (ASSR), also known as the envelope following response (EFR), is the auditory evoked potential elicited by an AM carrier (e.g., Purcell *et al.*, 2004) or a train of clicks/TBs (e.g., Mertes and Leek, 2016) presented at a given rate. In humans, ASSRs are typically measured from electrodes placed on the scalp, where common electrode sites include the high forehead, lower forehead, mastoid process, earlobe, and nape. Analysis of ASSR recordings typically aims to quantify the amplitude and phase synchrony of the responses at the modulation frequency. Clinical applications of the ASSR include screening for hearing loss in children (Cone-Wesson *et al.*, 2002), determining the presence of auditory function at levels beyond the upper limit for which ABRs are present (Vander Werff *et al.*, 2002), and evaluating auditory neuropathy (Rance *et al.*, 2005). In human research, ASSRs have been used to assess the possibility of cochlear synaptopathy (e.g., Bharadwaj *et al.*, 2015), the strength of the medial olivocochlear reflex by measuring suppression of the ASSR from contralateral noise (Mertes and Leek, 2016), and the neural circuits associated with schizophrenia (Thune *et al.*, 2016).

Recently, Chen and Jennings (2022) measured CAPs from an electrode on the tympanic membrane in eight normal-hearing human subjects and showed that an AM carrier elicits a CAP in response to each modulation cycle, resulting in a recording that resembles the ASSR. They designated this evoked potential “CAP_ENV_,” given that the response is characterized by a series of CAPs that follow the envelope of the stimulus.

Simulations of CAP_ENV_ where obtained using the stimuli from Chen and Jennings (2022), which consisted of a 500-ms, 3-kHz carrier that was sinusoidally amplitude modulated at 40, 80, 160, 320, 640, and 1000 Hz. The carrier level was 80 dB SPL. Simulated CAP_ENV_ waveforms were normalized according to the largest amplitude of the human-subject CAP_ENV_, which occurred for the first CAP in response to the 640-Hz modulation frequency and was 0.91 *μV*. Specifically, the un-normalized CAP_ENV_ model waveform for a given modulation frequency was multiplied by 0.91 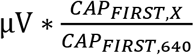, where CAP_FIRST,640_ is the amplitude of the first CAP of the simulated waveform for the 640 Hz modulation frequency, and CAP_FIRST,X_ is the amplitude of the first CAP of the simulated waveform for the modulation frequency of interest. Simulated CAP_ENV_ traces were compared with human CAP_ENV_ traces recorded from the average of seven normal-hearing adults from Chen and Jennings (2022). This comparison included expressing CAP_ENV_ in the time and frequency domains. Furthermore, contributions to CAP_ENV_ (80 Hz modulation frequency) from individual model ANFs were considered.

### B) Results and discussion

Fig. 10 presents the human-subject (Fig. 10 A) and simulated (Fig. 10 B: Zilany model, Fig. 10 C: Verhulst model) CAP_ENV_ waveforms for several modulation frequencies (rows). Human-subject and simulated CAP_ENV_ waveforms are qualitatively similar. That is, each waveform is characterized by a series of CAPs phase locked to the modulation frequency, where the largest CAP typically occurs at the onset of the stimulus. Furthermore, the CAP_ENV_ waveforms for modulation frequencies > 80 Hz exhibit adaptation, where the amplitudes of individual CAPs decrease over the duration of the stimulus. This adaptation occurs quickly (i.e., within the first 20 ms) for CAPs_ENV_ from human-subjects and those obtained with the Zilany model, whereas the adaptation is more gradual for responses obtained with the Verhulst model.

The top row of Fig. 11 presents the spectra of the human and simulated CAP_ENV_ waveforms for each modulation frequency. The spectra are qualitatively similar for CAP_ENV_ from human subjects and simulations. Specifically, spectral peaks occur at the modulation frequency and the harmonics of this frequency. The CAP_ENV_ waveforms from human subjects are associated with a higher noise floor than those from model simulations, which results in the absence of observable higher-frequency harmonics. The bottom row of Fig. 11 shows the spectral magnitude at the modulation frequency for human and simulated CAP_ENV_ waveforms in the form of a temporal modulation transfer function (Viemeister, 1979). The spectral magnitudes of the human CAP_ENV_ exhibit a shallow peak at 160 Hz. The spectra of the simulated CAP_ENV_ obtained with the Zilany and Verhulst are broadly consistent with this finding; however, the peak for the Verhulst model is located at a higher modulation frequency (640 Hz).

The contribution of individual model ANFs to the simulated CAP_ENV_ (3000 Hz AM carrier, 80 Hz modulation) is illustrated in Fig. 12 for the Zilany (Fig. 12A) and Verhulst (Fig. 12B) models. This figure is organized in the same format as Figs. 6 and 7, where the bottom traces display the PSTH from individual model ANFs, and the middle and top row plots show the PPSTH and CAP_ENV_, respectively. Phase locking to the modulation frequency is observed for all model ANFs; however, the depth of this modulation is greater for ANFs with CFs away from the carrier frequency (3000 Hz) compared to those near the carrier frequency (cf. the PSTH for CF=3.3 Hz versus that for other CFs). This result is consistent with synchrony capture, which is the finding that the response of ANFs with CFs tuned to formant frequencies is dominated by phase-locking to the frequency of the nearest harmonic (Deng and Geisler, 1987; Carney, 2018), rather than phase-locking to the modulation frequency (i.e., the fundamental frequency). For 100% modulation depth – as was the case for the current simulations – the carrier amplitude is larger than that of the side bands and therefore dominates the phase-locked activity of ANFs tuned near the carrier frequency. Thus, CAPs elicited by an AM carrier (i.e., CAP_ENV_) emerge primarily from stimulation of off-frequency ANFs according to the analysis in Fig. 12. Although CAP_ENV_ appears qualitatively similar for the Zilany and Verhulst model simulations, the contribution of high-CF ANFs to CAP_ENV_ is greater in the Verhulst model, while the Zilany model exhibits a relatively larger contribution from low-CF ANFs.

## VII. GENERAL DISCUSSION AND CONCLUSIONS

The objective of this study was to test the hypothesis that human CAPs elicited by a variety of acoustic stimuli could be effectively simulated by convolving a human-based UR with a PPSTH obtained from humanized versions of computational models of the auditory periphery. This hypothesis was supported by the ability of the simulations to *qualitatively* account for 1) the level-dependent amplitude, latency, and morphology of human CAPs elicited by acoustic clicks (Figs. 2 and 3), 2) a chirp benefit for chirp levels below 70 dB peSPL (Fig. 5 B,C), 3) the general trends in amplitude and latency for derived narrowband CAPs evoked by clicks and 2-kHz TBs (Fig. 8–9), and 4) the dependence of the CAP_ENV_ waveforms and spectra on modulation frequency (Figs. 10 and 11) for responses evoked by a 3-kHz sinusoidally AM carrier with modulation frequencies from 40 to 1000 Hz. These findings support the convolution framework (Goldstein and Kiang, 1958) as an effective approach for simulating the human CAPs. Moreover, the incorporation of AN activity from computational models, which account for cochlear nonlinearity and neural adaptation, provides a physiologically realistic prediction of the several level- and stimulus-dependent aspects of the human CAPs.

In some instances, simulated CAPs deviated from the human-subjects CAPs. These deviations are discussed for general trends observed for both models, as this study was not intended to provide a rigorous comparison between the Zilany and Verhulst models. Deviations are most apparent for simulations associated with chirps (Figs. 4 and 5). Although simulations from both models predicted a chirp benefit at low levels, the amplitude, latency, and morphology of simulated CAPs for chirp levels >70 dB peSPL were inconsistent with results from human subjects. Simulations for chirps >70 dB peSPL were associated with an early-latency component that was not observed in human-subject CAPs. Analysis of the contributions of individual ANFs to the chirp-evoked CAP (Fig. 7) indicated that this early-latency component is a result of upward spread of excitation of low-frequency components of the chirp. This observation suggests that some aspects of the upward spread of excitation of the human cochlea – which is determined by level-dependent cochlear tuning – are not captured by the humanized versions of the Zilany and Verhulst models. This observation is further supported by the findings that 1) simulations underpredicted the change in N_1_ latency for CAPs evoked by clicks (Fig. 3) (Verhulst et al., 2015), and 2) the range of latencies for derived narrowband responses is smaller for model simulations than for human-subject CAPs (Fig. 8). The foundation for human cochlear tuning for the Zilany and Verhulst models is based on OAE group delays obtained for low-level stimuli (Shera *et al.*, 2002). These OAE data define cochlear tuning for the models at low levels, but do not specify how this tuning changes with increasing sound level. As discussed by Verhulst *et al.* (2015), tuning at high stimulus levels is difficult to quantify using OAE techniques and that similar challenges are associated with psychophysical approaches as these depend strongly on the masking technique. Thus, guidance is currently lacking on how to model the details of level-dependent tuning of the human cochlea.

The development of computational models of the human peripheral auditory system have relied on animal experiments involving the measurements of basilar membrane motion and single-unit neural activity. The humanized versions of the Zilany and Verhulst models also rely on human-subject data from OAEs and evoked potentials sensitive to brainstem function (e.g., ABR, EFR). The simulations from the current study show how these models could use data from the human CAP, in addition to OAEs and other evoked potentials, to develop and refine predictions of responses from the human AN. Therefore, the approach described here could serve as a vehicle for identifying deviations between model simulations and empirically-measured CAPs, and those deviations could lead to a fruitful interplay between data and simulations that ultimately results in improved predictions of human AN responses.

The predicted CAPs presented in this paper are a first step in using humanized computational models to simulate the evaluation of the human cochlea using ECochG. These simulations were limited to the CAP and did not account for other cochlear potentials such as the CM (Chertoff *et al.*, 2012) or SP. Nevertheless, the approach presented herein is a useful tool for the design and analysis of experiments centered on measuring the human CAP. For example, this approach could be used to predict the contributions of individual ANFs to the human CAP elicited by stimuli beyond those studied in this paper (e.g., Lichtenhan *et al.*, 2013; Jennings and Dominguez, 2022), and novel stimuli associated with the design of a new experiment. Importantly, computational models often include options for simulating hearing loss by adjusting IHC and OHC health (Vecchi *et al.*, 2022) and simulating aging or synaptopathy by limiting the number of model ANFs (e.g., Lopez-Poveda and Barrios, 2013). Thus, the approach for simulating CAPs presented in this paper could be fruitful for designing experiments centered on investigating hearing loss, aging, and/or cochlear synaptopathy by comparing predicted CAP amplitudes – elicited by a given experimental stimulus – for normal and impaired model simulations.

## ACKNOWLEDGMENTS

This work was supported by grant K23 DC014752 from NIH/NIDCD (PI: Jennings) and by the University of Utah Undergraduate Research Opportunity Program (UROP). The authors thank Laurel Carney’s lab at the University of Rochester for the helpful comments on the work. The computations in this paper were performed using resources provided by the Center for High-Performance Computing at the University of Utah

